# Predicting metabolomic profiles from microbial composition through neural ordinary differential equations

**DOI:** 10.1101/2022.06.23.497381

**Authors:** Tong Wang, Xu-Wen Wang, Kathleen Lee-Sarwar, Augusto A. Litonjua, Scott T. Weiss, Yizhou Sun, Sergei Maslov, Yang-Yu Liu

## Abstract

Characterizing the metabolic profile of a microbial community is crucial for understanding its biological function and its impact on the host or environment. Metabolomics experiments directly measuring these profiles are difficult and expensive, while sequencing methods quantifying the species composition of microbial communities are well-developed and relatively cost-effective. Computational methods that are capable of predicting metabolomic profiles from microbial compositions can save considerable efforts needed for metabolomic profiling experimentally. Yet, despite existing efforts, we still lack a computational method with high prediction power, general applicability, and great interpretability. Here we develop a new method — mNODE (Metabolomic profile predictor using Neural Ordinary Differential Equations), based on a state-of-the-art family of deep neural network models. We show compelling evidence that mNODE outperforms existing methods in predicting the metabolomic profiles of human microbiomes and several environmental microbiomes. Moreover, in the case of human gut microbiomes, mNODE can naturally incorporate dietary information to further enhance the prediction of metabolomic profiles. Besides, susceptibility analysis of mNODE enables us to reveal microbe-metabolite interactions, which can be validated using both synthetic and real data. The presented results demonstrate that mNODE is a powerful tool to investigate the microbiome-diet-metabolome relationship, facilitating future research on precision nutrition.

## Introduction

Metabolic activities of microbial communities play an important role in shaping their biological functions and their interactions with hosts or environments. For example, a key way the human gut microbiome affects host physiology involves the production of small molecules [1, 2, 3]. Microbes in the gut can digest dietary fibers (i.e., undigestible carbohydrates like cellulose, hemicellulose, lignins, pectins, mucilages) [4] and eventually produce important metabolites such as essential amino acids [5] and short-chain fatty acids [2], which are crucial for human well-being [6, 7, 8]. Microbiota-derived metabolites, directly and indirectly, also influence the blood metabolites [9, 10]. It has been found that the set of metabolites in the human fecal metabolome has a significant overlap with metabolites detected in the blood [9]. To understand and eventually control microbial communities by modulation of nutrients or probiotic administration, the first step is to connect the metabolomic profile of a microbial community with its microbial composition.

Experimental measurement of metabolomic profiles usually requires metabolomics techniques such as LC-MS (Liquid Chromatography-Mass Spectrometry) or GC-MS (Gas Chromatography-Mass Spectrometry). However, metabolomics experiments are difficult because of expensive equipment [11, 12, 13], lack of automation [14, 15], and a limited metabolite coverage [11, 16]. In contrast, experimental measurements of microbial compositions through amplicon or shotgun metagenomic sequencing are less expensive, more easily automated, and provide good microbial coverage [17, 18]. As a result, compared to metabolomic data, amplicon or shotgun metagenomic sequencing data are more readily available for complex microbial communities. Hence, it is desirable to develop a computational method to predict metabolic profiles based on microbial compositions. Moreover, such a method could facilitate our understanding of the interplay between microorganisms and their metabolites.

Numerous computational methods have been proposed to achieve this goal, and they can be divided into the following three categories. (1) Reference-based methods such as MAMBO [19], MIMOSA [20], and Mangosteen [21]. In MAMBO (Metabolomic Analysis of Metagenomes using flux Balance analysis and Optimization), reference microbial genomes are used to reconstruct genome-scale metabolic models (GEMs). Then the metagenomic community abundance profile (i.e., the microbial composition) is correlated with the biomass production of the GEMs (obtained through the flux balance analysis). Finally, the correlation is optimized by multiple iterations of a Monte Carlo-based sampling algorithm. Both MIMOSA (Model-based Integration of Metabolite Observations and Species Abundances) and Mangosteen (Metagenome-based Metabolome Prediction) are reference-based, gene-to-metabolite prediction methods. Note that all those reference-based methods rely heavily on the completeness and accuracy of queried databases and GEMs. (2) Ecology-guided methods, where abundances of both microbes and metabolites are considered as end-products of the metabolic cascade, propagating through ecological networks of microbial communities (i.e., metabolite consumption and byproduct generation reactions by microbes) [22, 23, 24, 25]. Those methods heavily rely on the completeness and accuracy of ecological networks of microbial communities. (3) Machine learning (ML)-based methods, which are trained from paired microbiome and metabolome datasets, and then used to predict the metabolic profile of a never-seen microbiome sample based on its microbial composition, without using any reference database or domain knowledge regarding relationships between genes and metabolites. Various ML techniques such as elastic net [26], sparsified NED (Neural Encoder-Decoder) [27], multilayer perceptron [28], and word2vec [29] have been employed to predict metabolic profiles from microbial compositions. Despite the fact that these ML-based methods circumvent limitations of reference-based or ecology-guided methods discussed above and have been shown to generate promising results in various contexts, none of these ML-based methods utilize state-of-the-art deep neural network models such as the Neural Ordinary Differential Equation (NODE) [30], so their performance has not been fully maximized.

As a new family of deep neural network models, NODE has not been employed heavily in computational biology. Residual neural network (ResNet) is the precursor of NODE. As the most important characteristic of ResNet, the skip connection is highly analogous to the Euler step in ODE solvers [31, 32]. More specifically, one residual block can be regarded as a small time-change of variables in the language of ODEs [31, 32]. This analogy fostered the invention of NODE [30]. Rather than specifying a discrete sequence of hidden layers like ResNet, NODE parameterizes the derivative of the hidden state using a neural network with output computed using an ODE solver. In a sense, NODE represents continuous-depth models, has constant memory cost, adapts its evaluation strategy to each input, and can explicitly trade numerical precision for speed [30, 33]. As a result, higher accuracy can be achieved with fewer parameters and without explicitly introducing many neural layers [30, 33].

Here we leverage the power of NODE and propose a computational method called mNODE (Metabolomic profile predictor using Neural Ordinary Differential Equations) to predict metabolomic profiles from microbial compositions. We first generated synthetic data using the microbial consumer-resource model with cross-feeding interactions to validate that (1) mNODE outperforms existing ML-based methods, and (2) supplementing the information about nutrients as an additional input variable results in more accurate metabolome predictions, especially when the nutrient similarity across samples is low. We then compared mNODE with existing ML-based methods using microbiome and metabolome data collected in real microbial communities, finding that mNODE outperforms all existing ML-based methods. Moreover, we demonstrated that we could further improve the performance of mNODE by including food profiles from food frequency questionnaires as an additional input. In addition, to reveal microbe-metabolite interactions, we performed susceptibility analysis using mNODE to study how the predicted concentration of one metabolite responds to a perturbation in the relative abundance of one microbial species. We demonstrated clearly that susceptibilities for all microbe-metabolite pairs can be used to accurately predict microbe-metabolite interactions of synthetic data generated by the microbial consumer-resource model. Finally, we found genomic evidence strongly supports lots of predicted microbe-metabolite interactions from a human gut microbiome dataset.

## Results

### Overview of mNODE

We aim to predict the metabolomic profile of a microbial community based on its microbial composition and the potentially available information about diets. Fig. 1 shows a hypothetical example of two training scenarios when we follow the same model training protocol for mNODE: (1) without the dietary information as part of the input, and (2) with dietary information as part of the input. This demonstrative example system comprises 3 microbial species, 2 dietary items, 4 metabolites, and 15 samples. 10 samples are used in the training set and the remaining 5 samples are in the test set. We want to compare the performance of predicting metabolomic profiles with and without the dietary information in the input to understand to what extent the existence of dietary information helps our predictions. For a given set of samples with paired microbiome and metabolome data, we randomly split samples into two non-overlapping sets: the training set and the test set. We performed a 5-fold cross-validation on the training set to determine the optimal set of hyperparameters (i.e., the dimension for hidden layers and the weight parameter for the L2 regularization in mNODE) that maximizes the prediction power. Here, for each metabolite, we can calculate the Spearman’s Correlation Coefficient (SCC), denoted as *ρ*, between its predicted and experimentally observed concentrations across different samples. The overall prediction power of any computational method such as mNODE is evaluated as the mean SCC, 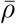, of all metabolites measured in experiments. After the 5-fold cross-validation, we retrained the model with the optimal set of hyperparameters on the entire training set and then used this trained model to generate the prediction on the test data (Fig. 1c).

**Figure 1:**
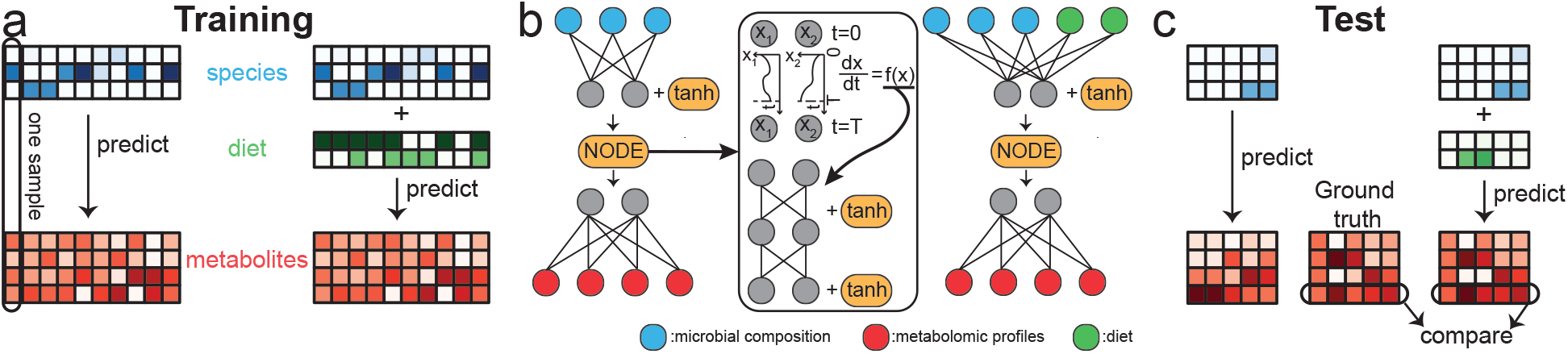
The mNODE workflow to predict metabolomic profiles from species microbial compositions and other dietary information. Across all panels, blue-colored arrays represent the microbial composition. Green-colored arrays represent the dietary information. Red-colored arrays represent the metabolomic profiles. None of the arrays has missing data and white cubes in all arrays just mean small values. 15 hypothetical samples in total with 3 species, 2 dietary items, and 4 metabolites are used to illustrate the idea. The samples are divided into training and test sets with the 2/1 ratio. **a** 10 samples in the training set are used for training machine learning models. There are two ways to predict metabolomic profiles: one without diets included in the input and the other one with diets included in addition to microbial compositions. **b** The architecture of mNODE for two training approaches. The neural ODE is a module in the middle of the architecture and it computes the time evolution of ODEs whose first-order time derivatives are approximated by an MLP with one hidden layer. Grey nodes represent neurons in hidden layers of mNODE. **c** Fully trained mNODE can generate predictions for metabolomic profiles in the test set. For each metabolite, the Spearman’s rank Correlation Coefficient *ρ* between its predicted profiles across samples and its true profiles in the test set is computed to quantify its predictability.

Since the numbers of microbial species (*N*_s_) and metabolites (*N*_m_) are generally different from each other, it is not possible to directly apply the original NODE (neural ODE) architecture [30], which requires the input and output dimensions to be equal. Here, we leverage the flexibility of the Multilayer Perceptron (MLP) in the data dimension and combine MLP with NODE to learn the mapping from inputs (microbial compositions) to outputs (metabolite concentrations). To be more specific, we introduce the NODE as a module in the middle, stuck between two densely connected layers (Fig. 1b). Our mNODE starts with one densely-connected layer (solid lines from blue nodes to grey nodes in Fig. 1b), followed by a non-linear activation function (the hyperbolic tangent function “tanh”). The densely-connected layer maps the input to the hidden layer of dimension *N*_h_. After passing through the densely-connected layer, the NODE module operates on the hidden space (neurons in hidden layers are denoted as greys nodes in Fig. 1b) of dimension *N*_h_. Mathematically, our NODE module maps the hidden variable 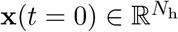 at time *t* = 0 to its state 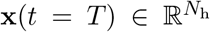 at time *t* = *T* following the ODE 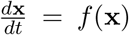 where *f*(x) is approximated by the learnable MLP with one hidden layer with dimension *N*_h_ (Fig. 1b, middle). Eventually, to generate the metabolome prediction from the hidden variable, we introduce another fully connected layer (solid lines from grey nodes to red nodes in Fig. 1b). In this study, we did not attempt to predict changes in metabolomic profiles over time; instead, the essential idea is to use NODE to approximate a function by modeling its first-order derivative via MLP with respect to continuous layers to reduce memory and training time. The L2 regularization is adopted to prevent overfitting. In total, there are two hyperparameters in mNODE to calibrate: the dimension of the hidden layer N_h_ and the weight parameter for L2 regularization λ.

### Validate mNODE using synthetic data

To validate mNODE, we generated synthetic data using the Microbial Consumer-Resource Model (MiCRM) which accounts for nutrient competition and cross-feeding interactions [34]. We created each independent synthetic community (“sample”) via sampling a set of species from a metapopulation pool and a set of nutrients from a nutrient pool to initiate the community assembly and then collecting the steady-state microbial species relative abundances as microbial compositions and steady-state nutrient concentrations as metabolomic profiles. Specifically, we assumed a metapopulation pool of 10 species and a nutrient pool of 10 nutrients with predetermined nutrient supply rates for all nutrients. To introduce the variability across samples, we assumed that each species from the metapopulation pool is introduced into each sample with the species sampling probability *p*_*s*_ and each nutrient from the nutrient pool is provided to the same ecosystem with its predetermined supply rate with the nutrient sampling probability *p*_n_. After the sampling event determines the initially introduced species and nutrients, the dynamics of species growth, nutrient consumption, and byproduct generation are simulated until the system reaches a steady state. The species relative abundances and nutrient concentrations in steady states from independent assembly simulations are collected as the microbial compositions and metabolomic profiles in the synthetic data (see Methods for details).

To explore whether mNODE generates better metabolome prediction than previously existing ML-based methods, we generated 300 samples with species sampling probability *p*_s_ = 0.5 and nutrient sampling probability *p*_n_ = 0.6. We left out 20% of all independent samples (60 samples) as the test dataset, whereas the remaining 240 samples are used as the training dataset. Figs. 2a1-a3 show the performance comparison between mNODE and previous ML-based methods (MelonnPan [26], sparse NED [27], MiMeNet [28], and ResNet [35]) through 3 metrics: (1) mean SCC, 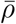 (Figs. 2a1, b1, c1), (2) the mean of the top-5 SCCs, 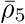 (Figs. 2a2, b2, c2), and (3) the number of metabolites (same as nutrients in the Microbial Consumer-Resource Model) with SCCs larger than 0.5, *N*_*ρ*>0.5_ (Figs. 2a3, b3, c3). The first performance metric was chosen to capture the overall performance, while the last two metrics were chosen to characterize the predictability of well-predicted metabolites and how many metabolites can be well-predicted respectively. We found that mNODE generates better predictions than all other methods in all three performance metrics. In addition, we intended to explore the usefulness of dietary information (i.e. nutrient supply rates) in improving the metabolome prediction of mNODE. This was achieved by incorporating nutrient supply rates as well as microbial composition as the input of mNODE (denoted as “mNODE + nutrients” in Figs. 2a1-a3). We found that incorporating nutrient information into mNODE offers the best prediction among all approaches among three metrics.

**Figure 2:**
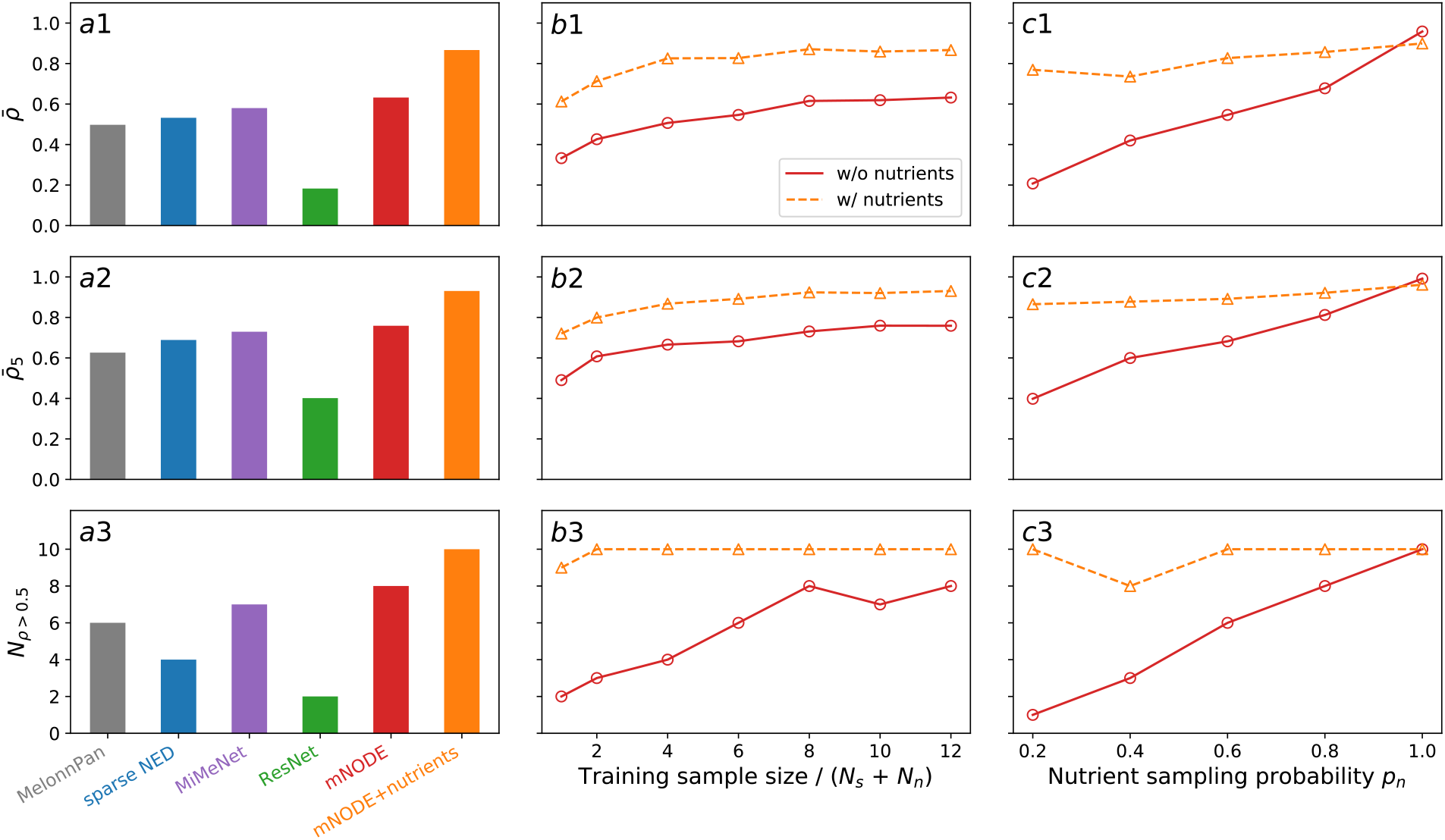
Model comparison and validation of mNODE on synthetic data generated by the microbial consumer-resource model. The predictive performance of mNODE is compared to other methods through 3 metrics: **a1** the mean SCC (Spearman Correlation Coefficient) 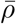, **a2** the top-5 mean SCC 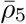, and **a3** the number of metabolites with an SCC larger than 0.5 *N*_*ρ*>0.5_. **b1-b3** The predictive performance of mNODE as a function of training sample size when the nutrient sampling probability (i.e. the fraction of nutrients externally supplied) *p*_n_ = 0.6 and the species sampling probability (i.e. the fraction of species introduced) *p*_s_ = 0.5. **c1-c3** The predictive performance of mNODE when the training sample size is 240 and the species sampling probability *p*_s_ = 0.5. Solid lines with circles are predicted results when nutrient supply rates (i.e. diets in the microbial consumer-resource model) are not included in the input of mNODE. Dashed lines with triangles are predicted results when nutrient supply rates are included in the input of mNODE.

The previous comparison utilized a fixed training sample size of 240. We would like to know how many samples are necessary to achieve the best performance for the system we investigated here. To guarantee the consistent measurement of model performance on the test set, we stuck to the same random train-test split used in Figs. 2a1-a3, reduced the training samples (through a subsampling of the original training set with 240 samples) while keeping the test set to be the same, trained the mNODE with fewer training samples, and measured 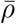 using the same test set of 60 samples. When nutrient supply rates are not used in the input, the performance reaches saturation when the training sample size reaches 8 times the sum of the number of nutrients *N*_r_(=10) and the number of microbial species *N*_s_(=10) shown by the saturating solid lines in Figs. 2b1-b3. In contrast, when nutrient supply rates are added to the input, the performance saturated as the training sample size is around 4 times of (*N*_r_ + *N*_s_) (saturating dashed lines in Figs. 2b1-b3), much lower than the training sample needed for the case without using nutrient supply rates. Additionally, it is worth noting that adding nutrient supply rates with a training sample size of 20 resulted in better performance than relying solely on the microbial compositions with a training sample size of 240 (Figs. 2b1-b3).

Besides the performance difference in the metabolome prediction between the two approaches caused by the training sample size, it is natural to speculate that the performance difference may also depend on the nutrient similarity across samples because the less variable nutrient information might render it less useful in discriminating metabolomic profiles. To test this, we systematically tune the nutrient sampling probability *p*_n_ because higher *p*_n_ implies more similar nutrient supplies. Figs. 2c1-c3 show how three performance metrics change as *p*_n_ increases. We found that when nutrient supply rates are not included in the input of mNODE, all 3 metrics increase as *p*_n_ increases from 20% to 100%: (1) 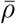 changes from 0.208 to 0.958; (2) 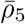 changes from 0.398 to 0.992; and (3) *N*_*ρ*>0.5_ changes from 0 to 10. When nutrient supply rates are included in the input, the performance of mNODE is consistently high over a wide range of values for *p*_n_. When *p*_n_ = 100%, there isn’t any variation in nutrient supply rates to independent communities across samples and thus nutrient supply rates become redundant, which explains why two approaches show similar performance (closer metric values in Figs. 2c1-c3).

### Superior performance of mNODE on real data

After the validation of mNODE using synthetic data, we systematically tested it using real data. A total of five datasets with paired microbiome and metabolome data were analyzed here. Those datasets were collected from microbial communities inhabiting diverse environments, such as human gut [36, 37, 38, 39, 40], human blood [37, 38, 39, 40], soil biocrust [41], and human lung [23]. Similar to the three metrics we used on synthetic data, here we assessed the model performance based on the mean SCC 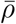, the top-50 mean SCC 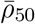, and the number of metabolites with an SCC larger than 0.5 (i.e., *N*_*ρ*>0.5_). We started with the dataset PRISM + NLIBD that collected fecal samples from CD (Crohn’s Disease) patients, UC (Ulcerative Colitis) patients, and healthy individuals [36]. This benchmark dataset is unique due to the presence of two IBD cohorts: a 155-member cohort collected at the Massachusetts General Hospital (PRISM) and a 65-member validation cohort collected in the Netherlands (NLIBD/LLDeep). Two cohorts were collected following the same protocols and processed in the same way (see Methods and Franzosa et al [36] for more details). For this dataset, we first selected hyperparameters for mNODE based on results of the 5-fold cross-validation of the PRISM cohort. Then, we trained mNODE with the selected hyperparameters on the entire PRISM cohort and computed predictions for the NLIBD cohort’s metabolome. Figs. 3a1-a3 show the performance of 5 methods on the test set NLIBD after their training process on PRISM. We found that mNODE had the highest 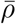 (0.294 compared to 0.191 from MiMeNet), the highest top-50 *ρ* (0.642 compared to 0.512 from MiMeNet), and the highest *N*_*ρ*>0.5_ (93 compared to 28 from MiMeNet) among all the ML-based methods. Furthermore, we directly compared *ρ* of annotated metabolites via a scatterplot in Supplementary Fig. S1. 76.8% of metabolites are better predicted by mNODE than the current start-of-the-art ML method: MiMeNet (Supplementary Fig. S1c). If we only focus on well-predicted metabolites (metabolites with *ρ* larger than 0.5 according to either mNODE or MiMeNet), 91.9% of metabolites are better predicted by mNODE than by MiMeNet (Supplementary Fig. S1d).

**Figure 3:**
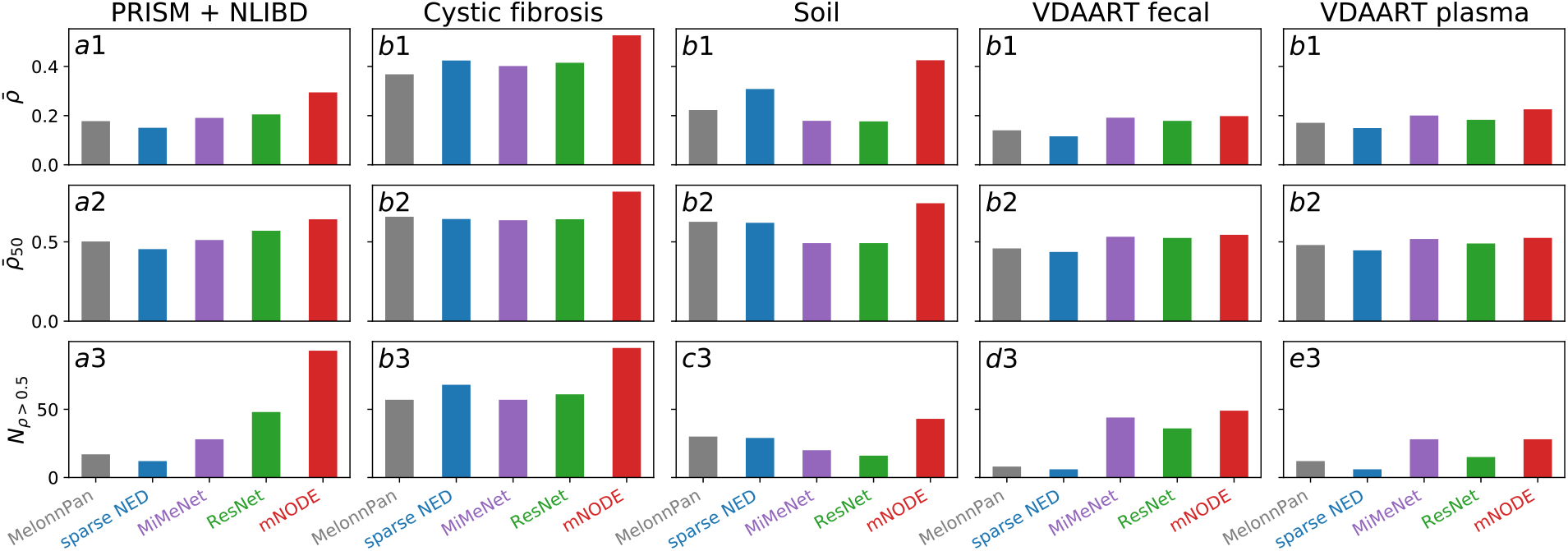
Performance comparison between mNODE and existing methods on real microbial community datasets. 3 metrics are adopted for comparing model performances: the mean SCC 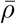, the top-50 mean SCC 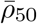, and the number of metabolites with SCCs larger than 0.5 *N*_*ρ*>0.5_. All datasets [36, 23, 41, 37, 38, 39, 40] are randomly divided into training and test sets with the 80/20 ratio except for the PRISM and NLIBD dataset [36]. **a1-a3** Performance of methods after training on the PRISM and test on the NLIBD [36]. **b1-b3** Performance of methods on the data from lung samples of patients with cystic fibrosis [23]. **c1-c3** Performance of methods on the data from soil biocrust samples after 5 wetting events [41]. **d1-d3** Performance of methods on the data from fecal samples of children at the age of 3 years [37, 38, 39, 40]. **e1-e3** Performance of methods on the data from blood plasma samples of children at the age of 3 years [37, 38, 39, 40].

To examine the superior performance of mNODE is independent of environments where microbial communities reside, we collected another 4 datasets: (1) lung sputum samples of 172 patients with cystic fibrosis [23], (2) 19 desert soil biocrust samples after the five continuous wetting events [41], (3) fecal samples of 340 children at the age of 3 years enrolled in the VDAART (Vitamin D Antenatal Asthma Reduction Trial) study [37, 38, 39, 40], and (4) blood plasma samples of the same 340 children in VDAART [37, 38, 39, 40]. The systematic comparison of different methods across the four datasets is shown in Fig. 3b-e. For lung samples from patients with cystic fibrosis, mNODE delivered the highest 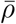 (0.526 compared to 0.402 from MiMeNet), the highest 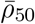 (0.817 compared to 0.636 from MiMeNet), and *N*_*ρ*>0.5_ (95 compared to 57 from MiMeNet). Regarding the desert soil biocrust samples, mNODE gave the highest 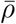 (0.425 compared to 0.178 from MiMeNet), the highest 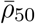 (0.743 compared to 0.492 from MiMeNet), and N_*ρ*>0.5_ (43 compared to 20 from MiMeNet). For fecal samples and plasma samples from children in VDAART, mNODE generated a much better prediction than MelonnPan [26] and sparse NED [27], and its predictive power is slightly better than MiMeNet (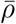 is 0.198 compared to 0.192 from MiMeNet for the fecal samples and 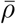 is 0.226 compared to 0.200 from MiMeNet for the plasma samples).

### Food profiles further improve the metabolome prediction

One of the unique features of the VDAART data is the documentation of food consumption frequency at the ages of 3 and 6 years. The food frequency questionnaire (FFQ) is commonly used to capture food and beverage consumption over time [42, 43, 44, 45]. The questionnaire consists of a finite list of foods and beverages with different choice answers that reveal the typical frequency of consumption over the specific time interval queried, such as the frequency of broccoli eaten in a week for the past year. FFQs come in many varieties, such as the Harvard Willett FFQ [42] and the NHANES (National Health and Nutrition Examination Survey) FFQ [43, 44, 45]. In VDAART, the FFQ followed and slightly modified the 87-item validated FFQ in preschool-age children [46]. Specifically, the FFQ in VDAART appears as a section in the 36 Months Quarterly Infant Follow-up Questionnaire [39]. Later, we converted their food consumption frequencies into the nutritional profiles based on the nutrient composition of each documented food. The conversion relies on the FNDDS (USDA’s Food and Nutrient Database for Dietary Studies) [47] database which encodes the detailed amounts of nutrient components in food and beverage items.

Equipped with the dietary information, we were ready to answer the question: which factor among the gut microbial composition, food consumption, and nutrient intake were most influential in determining the metabolomic profiles of children’ fecal samples and plasma samples? We addressed this question by training the mNODE in 5 different ways (foods profiles only, nutritional profiles only, microbial composition only, microbial composition with foods profiles, and microbial composition with nutritional profiles as the input of mNODE) and later comparing their differences in the prediction power. Among all approaches to predicting metabolomic profiles of either fecal or plasma samples, food profiles or nutritional profiles alone produced almost equally poor performance (Fig. 4). Adding nutritional profiles to the microbial composition as an extra input has little effect on its performance and its performance doesn’t show much difference from that of only using the microbial composition (Fig. 4). With regard to fecal samples, although 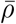 of metabolites predicted with the additional nutritional profiles (0.205) is a bit larger than 0.198 when nutritional profiles are not included, *N*_*ρ*>0.5_ with additional nutritional profiles is 41, less than 49 when they don’t belong to the input. Similarly, for plasma samples, 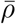 of metabolites predicted with nutritional profiles (0.219) is slightly smaller than 0.226 without nutritional profiles, *N*_*ρ*>0.5_ with nutritional profiles is 32, a bit larger than 28 when they are not present. The combination of microbial composition and food profiles seems to be the most successful approach, as indicated by the consistently best performance across 3 metrics and 2 data types (or environments). This shows that the variation of food consumption across individuals may help us to predict the metabolomic profiles better. One reason why the addition of food profiles may be more helpful than adding nutritional profiles is that the Bray-Curtis dissimilarity of nutritional profiles across paired samples is much lower than that of food profiles (Supplementary Fig. S2).

**Figure 4:**
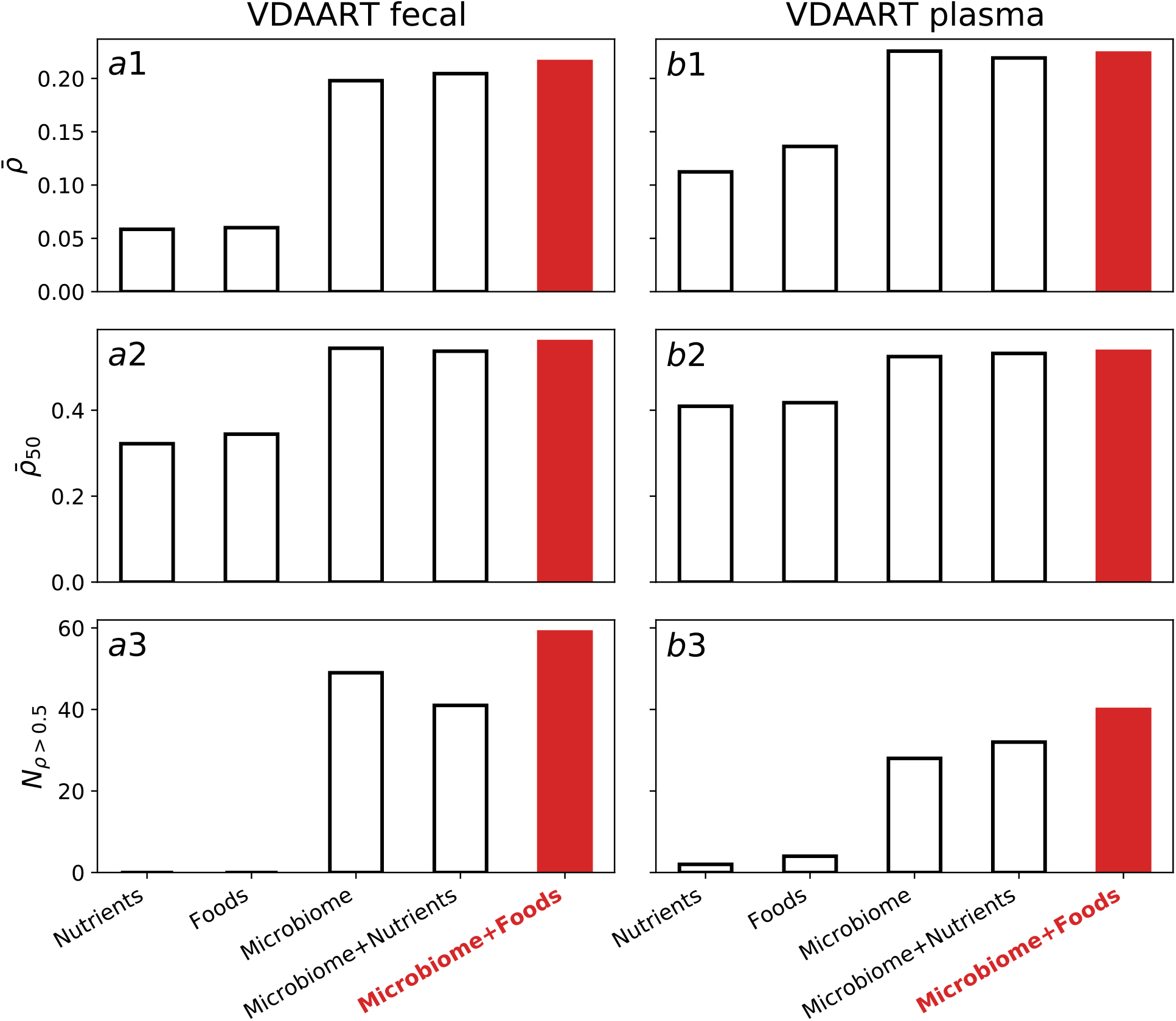
Performance of mNODE with different combinations of data types (microbial compositions, food profiles, and nutritional profiles) included in the input. 5 different combinations are used: foods only, nutritional profiles only, the microbial composition only, foods with microbial composition, and nutritional profiles with microbial composition. 3 metrics are adopted for comparing model performances: the mean SCC 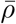, the top-50 mean SCC 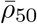, and the number of metabolites with SCCs larger than 0.5 *N*_*ρ*>0.5_. All datasets are randomly divided into the training set and test set with the 80/20 ratio. **a** Performance of methods on the data from fecal samples of children at the age of 3 years [37, 38, 39, 40]. **b** Performance of methods on the data from blood plasma samples of children at the age of 3 years [37, 38, 39, 40].

### Inferring microbe-metabolite interactions

Though deep learning models usually generate better predictions because they are better at approximating any functions, they still lack interpretability and hence offer little biological insights due to their “black-box” nature. To tackle this lack-of-interpretability issue, we proposed a perturbation method for mNODE to decipher the relationships between microbes and metabolites by measuring “susceptibilities” based on the response of metabolite concentrations to perturbation in inputs such as microbial relative abundances. Our method is similar to but simpler than the previously developed LIME (Locally Interpretable Model-agnostic Explanations) [48, 49]. To reveal the effect of species *i* on metabolite *α*, we perturbed the relative abundance of species *i* (*x*_*i*_) by a small amount Δ*x*_*i*_ for well-trained mNODE, re-predicted the concentration of metabolite *α*, and measured the deviation from the original prediction (we denoted the deviation as Δ*y*_*α*_). We defined the susceptibility of metabolite *α* to species *i* as 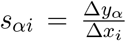 (Fig. 5a). The sign of *s*_*αi*_ reflects the influence of species *i* on metabolite *α*. For example, *s*_*αi*_ < 0 means that a higher abundance for species *i* leads to a lower predicted concentration for metabolite *α* and might imply that species *i* can consume metabolite *α*. Similarly, *s*_*αi*_ > 0 probably corresponds to the production of metabolite *α* by species *i*. More technical details about how to compute *s* _*ρ i*_ are in the Methods section.

**Figure 5:**
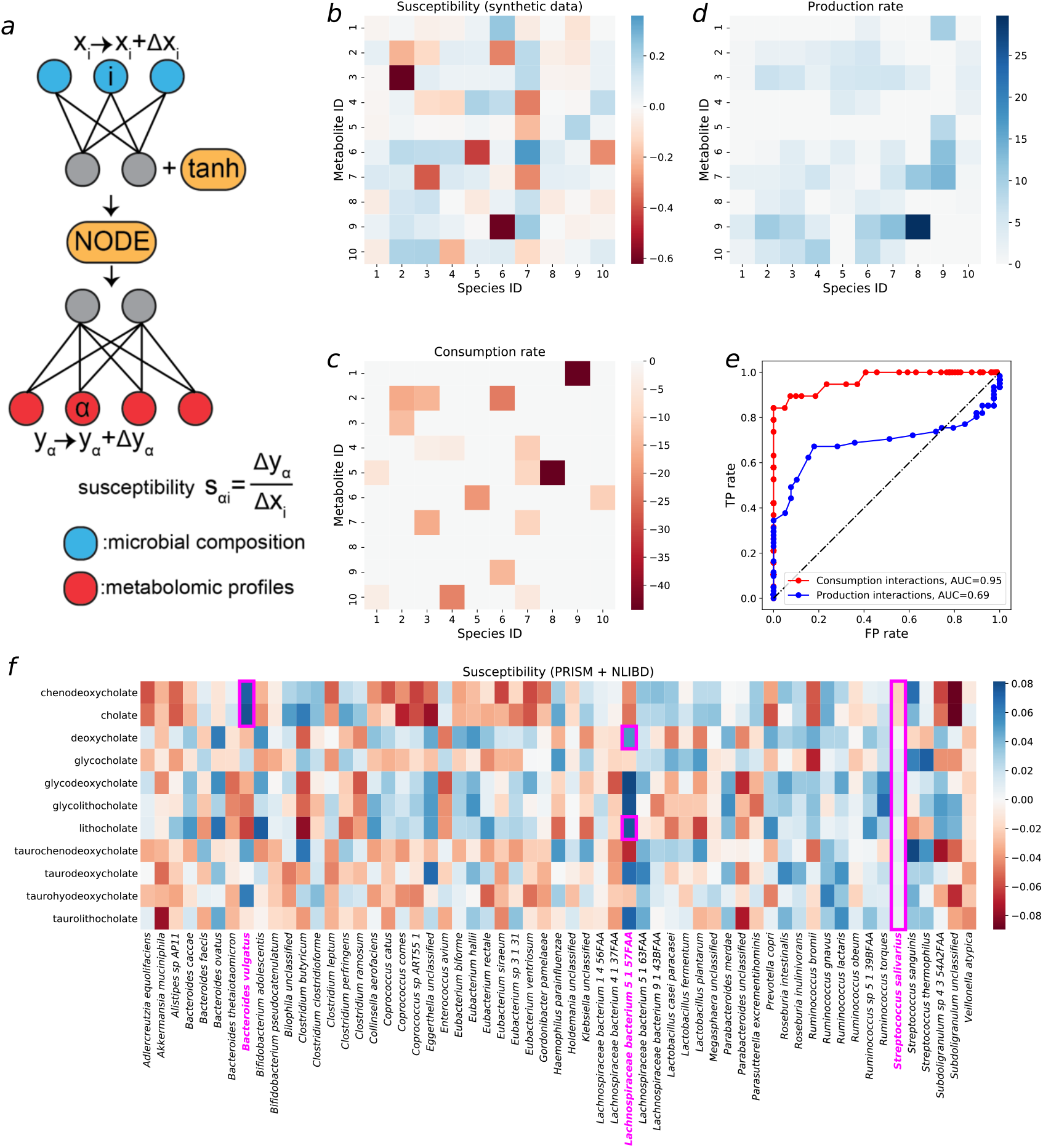
Using susceptibility of metabolite concentrations to microbial compositions of well-trained mNODE to infer microbe-metabolite interactions on both synthetic and real data (PRISM+NLIBD). **a** The susceptibility of the concentration of metabolite *α* (*y*_*α*_) to the relative abundance of species *i* (*x*_*i*_), denoted as *s*_*αi*_, is defined as the ratio between the deviation in the concentration of metabolite *α* (Δ*y*_*α*_) and the perturbation amount in the relative abundance of species *i* (Δ*x*_*i*_) **b** Susceptibility values for all microbe-metabolite pairs in the synthetic data used in Fig. 2a1-a3. **c** The ground-truth consumption matrix and corresponding rates in synthetic data. All consumption rates are shown as negative values for the convenience of comparison with panel b. **d** The ground-truth production matrix and corresponding rates in synthetic data. Details about how the production matrix is obtained can be found in the supplemental information. **e** The ROC (Receiver Operating Characteristic) curve based on TP rates and FP rates which are obtained by setting different susceptibility thresholds for classifications of interactions. **f** The susceptibility values for all microbe-metabolite pairs in PRISM + NLIBD [36] used in Fig. 3a1-a3.

To validate if the proposed susceptibility measure can decipher microbe-metabolite consumption and production interactions, we first tested the idea on the synthetic data used in Figs. 2a1-a3 and computed susceptibilities for all microbe-metabolite pairs (Fig. 5b). Synthetic data are well suited to validate the susceptibility method because it is possible for us to directly compare the susceptibility matrix (Fig. 5b) with the ground-truth consumption interactions (Fig. 5c) and production interactions (Fig. 5d) in the microbial consumer-resource model. It is visually evident that many ground-truth interactions in Fig. 5c are correctly revealed in the susceptibility matrix (indicated by having red colors in Fig. 5b). To systematically compare the susceptibility matrix with ground-truth consumption interactions, we classified all microbe-metabolite pairs into (1) having consumption interactions or (2) not having consumption interactions according to a classification threshold *s*_thres_. Only microbe-metabolite pairs that have susceptibilities lower than *s*_thres_ would be considered as predicted consumption interactions. Lowering such the threshold *s*_thres_ would only consider microbe-metabolite pairs with relatively small susceptibilities (i.e. negative values far away from zero) to be predicted as consumption interactions. We incrementally increased the threshold *s*_thres_ from the minimal value of susceptibilities to the maximal value, performed the classification task for each threshold, computed TP (True-Positive) rates and FP (False-Positive) rates for all thresholds, and measured the AUC (Area Under Curve) for the ROC (Receiver Operating Characteristic) curve. The same classification procedure can be applied to infer production interactions where only large susceptibility values would be classified as production interactions. We found that, for synthetic data, the susceptibility can predict consumption interactions with great performance (AUC = 0.95) and production interactions with decent performance (AUC = 0.69). To understand if the less accurate predictions for production interactions are due to their overlaps with consumption interactions (i.e. a metabolite can be consumed and produced by a species simultaneously; see Methods for more details), we designed a MiCRM (Microbial Consumer-Resource Model) with species-specific byproduct generation and no overlap between consumption and production interactions (more details in supplemental information). When production interactions don’t overlap with consumption interactions, the prediction power for production interactions is almost equivalent to that for consumption interactions (Fig. S3e).

We eventually applied the validated susceptibility method to all real-life datasets and susceptibility values for all datasets are provided as supplemental data/tables. Here we focused on the dataset PRISM + NLIBD because gut microbiomes of human adults have been investigated more extensively. For this purpose, we focused on the bile acid metabolism which is relatively well-studied [50, 51, 52, 53, 54] and is shown to be associated with human gastrointestinal diseases such as gastrointestinal cancers [53] and IBD [55, 56]. We carefully chose metabolites that are related to the primary, conjugated, and secondary forms of cholate and chenodeoxycholate. Heinken et al performed a systematic assessment of bile acid metabolism for more than 800 gut microbial strains based on their genomes [54]. In order to test if inferred interactions based on our susceptibility matrix have genomic evidence, we searched for some microbe-metabolite pairs with large absolute values for susceptibilities. *Bacteroides vulgatus* ATCC 8482 has large positive susceptibilities for cholate and chenodeoxycholate. This is supported by the genomic evidence that *Bacteroides vulgatus* ATCC 8482 contains the bsh (bile salt hydrolase) gene, which encodes the BSH enzyme responsible for the deconjugation of conjugated primary bile acids to primary bile acids cholate and chenodeoxycholate [50, 54]. Similarly, *Lachnospiraceae bacterium* 5 1 57FAA has large positive susceptibilities for deoxycholate and lithocholate, as evidenced by the presence of the bai pathway in its genome, which transforms primary bile acids cholate and chenodeoxycholate to secondary bile acids deoxycholate and lithocholate respectively [54]. It is worth noting that other strains of the species *Lachnospiraceae bacterium* don’t have large susceptibility values for deoxycholate and lithocholate, validated by the genomic evidence of the bai pathway not existing in genomes of *Lachnospiraceae bacterium* 5 1 63FAA and *Lachnospiraceae bacterium* 8 1 57FAA [54]. Susceptibility values close to zero are likely to imply no interactions between microbes and metabolites. For example, all susceptibilities of *Streptococcus salivarius* are much closer to zero compared with other entries in the susceptibility matrix and its genome indeed contains no genes related to bile acid metabolism [54].

## Discussion

Understanding the function of microbial communities requires the profiling of their metabolic activities. Metabolomics such as LC-MS applied to many microbial communities often profiles the end-product of microbial metabolism; metabolites not completely consumed and metabolites further produced by microbes together contribute to the metabolome that can be measured in the untargeted metabolomics. Metabolomics is much more expensive than metagenomics or 16S sequencing due to pricy equipment and laborious manual curation. Computational methods, capable of predicting metabolite levels using microbial composition, offer a much more cost-effective way to quantify the distribution of metabolite concentrations across samples. Moreover, such a model might help us better understand the interaction of microbes and metabolites. In this study, we developed the mNODE method, which produces better metabolome predictions than almost all previously developed methods. Moreover, we further improved the performance of mNODE by including food profiles together with the microbial composition as the input of mNODE. Finally, we showed that mNODE infers microbe-metabolite interactions accurately on synthetic data and a human dataset (PRISM+NLIBD) via susceptibility analysis. In the future, if more omics data such as the metatranscriptome and metaproteome are available, they can also be easily incorporated as inputs of mNODE to investigate the relative importance of each data type in predicting the metabolome.

NODE integrates the well-developed ODE solving techniques of the past 120 years with deep learning by “unrolling” the implicit layer that approximates first-order time derivatives of the dynamical systems. Because of this, researchers are increasingly interested in the possibility of solving dynamical systems problems using the architecture of NODE. For instance, recently Dutta et al showed that NODE can generate the solutions for the various evolution problems of different fluid dynamics and further provide a promising potential to extrapolate [33]. Another attempt was to use NODE to learn ecological and evolutionary processes based on the time-series data generated by traditional ecological models [57]. Microbial communities are dynamical systems, in which microbes interact primarily through the metabolite consumption and production of byproduct metabolites [58, 34]. nutrients provided periodically (such as dietary fibers in the diet for gut microbiomes) or continuously (such as nutrient fluxes in chemostats) to a microbial community can be consumed by microbes and converted to other byproducts by the microbes’ metabolism. The experimentally measured metagenome and metabolome of samples from microbial communities at different times can be considered as the profiling of microbial abundances and metabolites concentrations at corresponding times. Consequently, we expect that leveraging the NODE framework with the correct input data types to determine the community dynamics of microbes and their metabolism would generate better performance. Out of many input data types, we believe that diet/nutrient information is an important one.

However, the current method of documenting diet information via the FFQ has some limitations [59]. On one hand, the systematic error in the consumption amount occurs because the FFQ only surveys the frequency of eating one food item, rather than the amount of each food item eaten per meal. Certainly, it is challenging for the FFQ to document the precise food consumption information for each meal, as it usually enquires about the habitual diet over a long period of time such as a week, month, or year. Digital documentation of the amount of food eaten for every meal may alleviate this error [59]. On the other hand, the FFQ lacks detailed information about food product brands and food preparation. A similar issue arises when food profiles in FFQs are converted into nutritional profiles due to limited nutrient items and the lack of detailed nutrient composition. For example, dietary fiber in the conversion database encompasses all types of fibers. But different microbes have different consumption abilities and rates towards different fibers and polysaccharides [60, 61]. A detailed metabolomic analysis of a comprehensive set of foods might be important for the advancement of precision nutrition [62].

We notice that the prediction accuracy for microbial communities collected from lungs [23] and soils [41] is higher than that for human gut microbiomes (i.e. PRISM + NLIBD [36] and VDAART [37, 38, 39, 40]), regardless of computational approaches utilized. The mean Spearman Correlation Coefficient 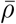 for lung samples (0.526 for mNODE and 0.402 for MiMeNet) is much higher than that for the VDAART fecal samples (0.198 for mNODE and 0.192 for MiMeNet). This demonstrates that metabolomic profiles of some microbiomes are harder to be predicted than others and we suspect that this is due to the difference in the ratio between the input dimension (e.g., the number of microbial taxa) and the output dimension (e.g., the number of metabolites in metabolomic profiles). For instance, for lung samples [23], the number of microbial taxa (1119) is much larger than the number of metabolites (168). As a comparison, for fecal samples in PRISM + NLIBD, the number of microbial taxa (200) is much smaller than the number of metabolites (8848). This comparison just showed the complexity of inferring metabolomic profiles for human gut microbiomes. Two datasets for human gut microbiomes still have different prediction performances; the mean Spearman Correlation Coefficient 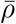 for PRISM + NLIBD (0.294 for mNODE) is slightly higher than that for the VDAART fecal samples (0.198 for mNODE). We believe that this may due to the fact that children investigated in VDAART probably don’t have well-established gut microbiomes as adults in PRISM + NLIBD.

It is worth noting that our mNODE method is very generic, which makes it easy to apply it to other biological problems where the dynamics of the system play an essential role. For example, it is possible to apply this model framework to predict one omics data type such as the metatranscriptome and metaproteome from the other omics data such as the metagenome. For many bacteria like *Staphylococcus aureus* and *Bacillus subtilis*, the expression of genes in the DNA to mRNAs and its further translation to proteins are dynamic processes that are tightly regulated [63]. Since the protein levels reflect metabolic activities of microbes better than their genomic information, if we can design a model to predict metaproteome from metagenome, we might be one step closer to unraveling the function of microbes in their communities. Also, such a connection between microbial genome and their proteome may enable us to predict phenotypes of microbes such as their specific growth rates in different environments if environmental data is available.

## Methods

### Datasets

#### PRISM + NLIBD fecal samples

This dataset was reported by a study related to the fecal microbiome and metabolome samples of IBD patients [36]. There are two cohorts involved in the study. The first cross-sectional cohort of individuals was collected by a study PRISM (the Prospective Registry in IBD Study at MGH). This cohort includes 68 patients with Crohn’s disease (CD), 53 patients with ulcerative colitis (UC), and 34 healthy controls. The second study cohort (NLIBD/LLDeep) was independently collected in the Netherlands and consists of 20 CD patients, 23 UC patients, and 22 healthy controls. Fecal samples from both cohorts were collected following the same protocol and later both the microbiome and metabolome are profiled and analyzed in the same way. In total, 200 microbial taxa were generated using the metagenomic shotgun sequencing and 8848 unique metabolites were obtained by four LC-MS (Liquid Chromatography-Mass Spectrometry) metabolomics methods. Out of 8848 unique metabolites, 466 are annotated. In all machine learning tasks, all data (microbial composition and metabolomic profiles) related to individuals in PRISM are used as the training set and the external validation cohort NLIBD/LLDeep is used as the test set.

#### Cystic fibrosis lung samples

All lung sputum samples from 172 patients were collected in a study that investigates how the chemical gradient drives the shift in the microbiome structure and the pattern of metabolite production [23]. 1119 microbial features were determined by the 16S rRNA gene sequencing and profiles of 168 metabolites were generated by the tandem liquid chromatography-mass spectrometry (LC-MS/MS) and gas chromatography-mass spectrometry (GC-MS) metabolomics. For all machine learning tasks, an 80/20 train-test split with the random state set as 42 is utilized to guarantee a fair comparison.

#### Desert soil biocrust samples

Swenson et al describes how biocrusts from four successional stages were wet up and sampled at five time points for each stage [41]. 466 dominant taxa were determined by the shotgun metagenomic sequencing that measures their single-copy gene markers. The liquid chromatography-mass spectrometry (LC/MS) soil metabolomics identified 85 metabolites that changed at least two-fold across both wetting and successional stages. An 80/20 train-test split with the random state set as 42 is utilized for all models.

#### VDAART children’s fecal samples, plasma samples, and FFQ data

VDAART (Vitamin D Antenatal Asthma Reduction Trial) is a trial of prenatal vitamin D supplementation to prevent asthma and wheezing offspring [37, 38]. Fecal samples and blood plasma samples of 340 children at the age of 3 years were collected. The 16S rRNA gene sequencing identified the microbiome profiles of 209 microbial taxa in fecal samples [40]. Primer and adapter trimming was performed using Skewer. Chimera checking and filtering were performed using Qiime2 [64]. Reads were denoised using DADA2 as implemented in Qiime2 [65]. The metabolomic profiles of both fecal and blood samples are generated via the using ultra-high performance liquid chromatography-mass spectrometry (UPLC-LC/MS) as previously described [66, 40, 39]. Identification of known chemical entities was based on a comparison to metabolomic library entries of purified standards based on chromatographic properties and mass spectra. 1298 metabolites were determined in the fecal samples and 1064 metabolites were determined in the blood samples. Besides, we extracted their diet information from the section in the 36 Months Quarterly Infant Follow-up Questionnaire that documented the food frequency consumed by children [39]. Child diet was evaluated at age of 3 years when parents completed a modified version of a semi-quantitative 87-item food frequency questionnaire (FFQ) that was previously validated in preschool-age children [46]. The food frequency table can be further converted to the nutritional profiles using FNDDS (USDA’s Food and Nutrient Database for Dietary Studies) which is a database that is used to convert food and beverages consumed into gram amounts and to determine their nutrient values. All 340 children are divided into either the train set or the test set with an 80/20 ratio with the random state set as 42.

### Mnode

The NODE (Neural Ordinary Differential Equations) is a deep learning method that combines explicit layers with implicit layers where the states of hidden layers are described by Ordinary Differential Equations. Our mNODE introduces the NODE as a module in the middle.

- Data processing: The CLR (Centered Log-Ratio) transformation is applied to microbial abundances and metabolite concentrations.
- Model detail: The architecture consists of 3 connected modules: (1) one fully connected layer that maps the input (such as the microbial composition) to the hidden layer with dimension *N*_h_ followed by an activation function, (2) Neural ODE module [30] where the first-order time derivative is approximated by a one-hidden-layer MLP with the hidden layer dimension the same as *N*_h_, and (3) one fully connected layer that maps from the hidden layers to the output (i.e., the metabolomic profiles). The L2 regularization with the weight parameter λ is assumed to prevent overfitting.
- Training method: Adam optimizer [67] is used for the gradient descent. The training stops if the loss on the validation/test set starts to increase within the past 20 epochs. The criterion for the increase is judged by whether the number of increases of the loss in the past 20 epochs (i.e., the loss at epoch i minus the loss at epoch i-1 is larger than 0) is larger than 12.
- Activation function: the hyperbolic tangent function tanh.
- Hyperparameter selection: Two hyperparameters are selected based on the 5-fold cross-validation results (the mean Spearman’s rank correlation coefficients) on the training set: the dimension of the hidden layer *N*_h_ and the L2 penalty with weight parameter λ. *N*_h_ is chosen from [32, 64, 128], and λ is selected from [10^*-*4^, 10^*-*3^, 10^*-*2^].

### Inferring microbe-metabolite interactions via susceptibility

The well-trained mNODE takes relative abundances of all species (relative abundance of species *i* is denoted as *x*_*i*_) as inputs and generate predictions for metabolite concentrations (concentration of metabolite *α* is written as *y*_*α*_). For the sample m in the training set, 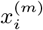 for all *i* is provided as the input vector to the trained mNODE and mNODE can predict concentrations 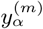 for all metabolites. To investigate the influence of species *i* on metabolite *α* for a particular sample *m*, we perturb 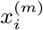 by setting it as the mean of relative abundances for species *i* across samples 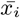 while keeping values of 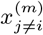 intact. Thus the perturbation amount for species *i* is 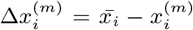. This perturbed vector for microbial relative abundances is provided to the trained mNODE to regenerate predictions for metabolite concentration 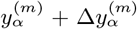 where 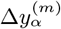 is the deviation of newly predicted concentration for metabolite *α* when mNODE uses the perturbed input vector from that when mNODE uses the unperturbed input vector. For the sample *m*, the susceptibility of metabolite *α* to species *i* can be defined as 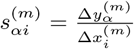. To properly take into account all training samples, we average susceptibility over samples and obtain the overall susceptibility of metabolite *α* to species *i*, 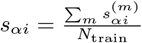, where *N*_train_ is the number of samples in the training set.

## Statistics

To calculate correlation coefficients throughout the study, we used Spearman’s rank correlation coefficient. Wherever we used *P* values, we explained in the Methods how we calculated them, since for all such measurements in the study, we calculated the associated null distributions from scratch. All statistical tests were performed using standard numerical and scientific computing libraries in the Python programming language (version 3.7.3), the Julia programming language (version 1.6.2) and Jupyter Notebook (version 6.1).

## Data and code availability

All code for simulations used in this manuscript can be found at https://github.com/wt1005203/mNODE.

## Acknowledgements

Y.-Y.L. acknowledges grants from the National Institutes of Health (R01AI141529, R01HD093761, RF1AG067744, UH3OD023268, U19AI095219, and U01HL089856). We thank Nancy Laranjo for VDAART data support.

## Author contributions

Y.-Y.L. conceived the project. T.W. and Y.-Y.L. designed the project. T.W. performed all the numerical calculations and data analysis. T.W. processed the real data with assistance from X.-W.W. and K.L.S. All authors analyzed the results. T.W. and Y.Y.L. wrote the manuscript. All authors edited and approved the manuscript.

