## Supplemental Information for "Predicting metabolomic profiles from microbial composition through neural ordinary differential equations"

### Supplementary Information

#### Existing machine learning methods

*MelonnPan.* MelonnPan is a computational method based on the regularized linear regression for predicting metabolite composition from microbiome sequencing data [26].

- Data processing: It applies the rank-transformation to the normalized microbial abundances. Specifically, it utilizes the quantile transformation that transforms features to follow a normal distribution. Normalized metabolite profiles are arcsine square root transformed as the outputs to be predicted by the model.
- Model detail: The model is a linear regression model with elastic net regularization. The elastic net regularization is a hybrid method that linearly combines the L1 (i.e., LASSO) and L2 (i.e., ridge) penalties. It has the elastic net mixing parameter $\alpha$ (the fraction of L2 between L1 and L2) to adjust the ratio between two types of regularizations and sparsity parameter $\lambda$ (penalty weight) to modify the overall parameter sparsity. For each metabolite, one elastic net model is designed to achieve the best performance.
- Hyperparameter selection: Two hyperparameters are selected based on the 5-fold cross-validation results (the mean Spearman's rank correlation coefficients) on the training set: the elastic net mixing parameter $\alpha$ and sparsity parameter $\lambda$. $\alpha$ is selected from [0.1, 0.5, 0.9] and $\lambda$ is selected from [$10^{-4}$, $10^{-3}$, $10^{-2}$, $10^{-1}$].

*Sparse NED.* Le et al proposed an MLP (Multiple-Layer Perceptron) model with one hidden layer with fewer model parameters to predict metabolite clusters [27].

- Data processing: It applies the CLR (Centered Log-Ratio) transformation [68] to both microbial abundances and metabolite concentrations since both omics data are compositional. After that, the Z-score transformation, widely used in machine learning, is applied to both data types.
- Model detail: The model is a sparsified MLP model with one hidden layer. the dimension of the hidden layer is $N_{\mathrm{h}}$. The learning process is made of two steps: the screening stage and the training stage. During the screening stage, the MLP model with fully connected weights is trained and connections that are most useful in extracting the information needed to predict metabolite concentrations from microbe abundances are identified. The connection importance is measured by the normalized magnitude of the derivatives. The connection with a larger magnitude for its derivative is considered to be more important. Only connections ranked as the top $\beta$ percentile are kept and $\beta$ is a sparsity parameter that can be adjusted to change the percent of weights kept. Then in the training stage, the MLP model with less important connection deactivated (or masked) from the forward-feed and backpropagation operations.
- Training method: Adam (Adaptive Moment Estimation) optimizer [67] is used for the gradient descent. the training stops after 100 epochs.
- Activation function: the hyperbolic tangent function tanh.
- Hyperparameter selection: Two hyperparameters are selected based on the 5-fold cross-validation results (the mean Spearman's rank correlation coefficients) on the training set: the dimension of the hidden layer

$N_{\mathrm{h}}$ and sparsity parameter $\beta$. $N_{\mathrm{h}}$ is selected from [32, 64, 128] and $\beta$ is selected from [0.05, 0.1, 0.2, 0.5].

*MiMeNet.* MiMeNet (Microbiome-Metabolome Network) is a method based on the multilayer perceptron neural network (MLPNN) with two ways to prevent the overfitting: L2 regularization and dropout [28]. It attempted to predict the entire metabolite profile and learn the microbe-metabolite interactions using the feature attribution scores [28].

- Data processing: The Z-score transformation is applied to the CLR (Centered Log-Ratio) transformed microbial abundances and metabolite concentrations.
- Model detail: The model is an MLP model with one hidden layer or several hidden layers. The L2 regularization with weight parameter $\lambda$ is adopted to sparsify the number of model parameters. In addition, the dropout with a rate $r$ at each hidden layer (i.e., a random fraction of nodes and corresponding weights are masked temporarily) is applied to further regularize the MLP model.
- Training method: Adam optimizer [67] is used for the gradient descent. The training stops if the loss on the validation/test set has not improved within the past 40 epochs.
- Activation function: Rectified Linear Unit (ReLU).
- Hyperparameter selection: Four hyperparameters are selected based on the 5-fold cross-validation results (the mean Spearman's rank correlation coefficients) on the training set: the dimension of the hidden layer $N_{\mathrm{h}}$, the number of hidden layers $N_{\ell}$, the L2 penalty with weight parameter $\lambda$, and the dropout rate $r$. $N_{\mathrm{h}}$ is selected from [32, 128, 512], $N_{\ell}$ is selected from [1, 2, 3], $\lambda$ is selected from [$10^{-4}$, $10^{-3}$, $10^{-2}$, $10^{-1}$], and $r$ is selected from [0.1, 0.3, 0.5].

*ResNet.* The ResNet (Residual neural Network) is a deep learning method based on the idea of residual blocks and skip connections.

- Data processing: The Z-score transformation is applied to the CLR (Centered Log-Ratio) transformed microbial abundances and metabolite concentrations.
- Model detail: We adapted the ResNet (Residual neural Networks) architecture from Goyal et al [35]. Goyal et al proposed the Linear-QuadraticResidual Network (LQResNet), which linearly combines the linear and quadratic mappings with the residual neural network to model the first-order time-derivative of variables [35]. The architecture consists of 3 connected modules: (1) one fully connected layer that maps the input (such as the microbial composition) to the hidden layer with dimension $N_{\mathrm{h}}$ followed by an activation function, (2) N residual blocks with each block made of a one-hidden-layer MLP with the layer dimension the same as $N_{\mathrm{h}}$ plus the skip connection, and (3) one fully connected layer that maps from the hidden layers to the output (i.e., the metabolomic profiles). The L2 regularization with the weight parameter $\lambda$ is assumed to prevent overfitting.
- Training method: RAdam (Rectified Adam) optimizer [69], which utilizes a warm-up strategy to rectify the variance of the adaptive learning rate, is used for the gradient descent. The training stops after 100 epochs.

- Activation function: Exponential Linear Unit (ELU).
- Hyperparameter selection: Three hyperparameters are selected based on the 5-fold cross-validation results (the mean Spearman's rank correlation coefficients) on the training set: the dimension of the hidden layer $N_\mathrm{h}$, the number of residual blocks $N$, and the L2 penalty with weight parameter $\lambda$. $N_\mathrm{h}$ is chosen to be the same as, 2 times, or 3 times the input dimension, $N$ is selected from [2, 3, 4], and $\lambda$ is selected from [1, 5, 20].

#### mNODE

*mNODE.* The NODE (Neural Ordinary Differential Equations) is a deep learning method that combines explicit layers with implicit layers where the states of hidden layers are described by Ordinary Differential Equations. Our mNODE introduces the NODE as a module in the middle.

- Data processing: The CLR (Centered Log-Ratio) transformation is applied to microbial abundances and metabolite concentrations.
- Model detail: The architecture consists of 3 connected modules: (1) one fully connected layer that maps the input (such as the microbial composition) to the hidden layer with dimension $N_\mathrm{h}$ followed by an activation function, (2) Neural ODE module [30] where the first-order time derivative is approximated by a one-hidden-layer MLP with the hidden layer dimension the same as $N_\mathrm{h}$, and (3) one fully connected layer that maps from the hidden layers to the output (i.e., the metabolomic profiles). The L2 regularization with the weight parameter $\lambda$ is assumed to prevent overfitting.
- Training method: Adam optimizer [67] is used for the gradient descent. The training stops if the loss on the validation/test set starts to increase within the past 20 epochs. The criterion for the increase is judged by whether the number of increases of the loss in the past 20 epochs (i.e., the loss at epoch i minus the loss at epoch i-1 is larger than 0) is larger than 12.
- Activation function: the hyperbolic tangent function tanh.
- Hyperparameter selection: Two hyperparameters are selected based on the 5-fold cross-validation results (the mean Spearman's rank correlation coefficients) on the training set: the dimension of the hidden layer $N_\mathrm{h}$ and the L2 penalty with weight parameter $\lambda$. $N_\mathrm{h}$ is chosen from [32, 64, 128], and $\lambda$ is selected from [$10^{-4}$, $10^{-3}$, $10^{-2}$].

#### Microbial Consumer-Resource Model (MiCRM) with cross-feeding interactions

Similar to the formalism of ecological models developed for microbial communities [34], we considered how the nutrients supplied to a microbial ecosystem are consumed and other byproduct nutrients are further produced by microbes. The supply rate of nutrient $\alpha$ is $h_\alpha$. Also, the system is assumed to be constantly diluted with a dilution rate $\delta$. For each microbial species $i$, the consumption rate of nutrient $\alpha$ per microbial density per nutrient concentration is assumed to be $a_{i\alpha}$. As a result, the overall consumption rate of nutrient $\alpha$ by species $i$ is $J^{con}_{i\alpha} = a_{i\alpha}N_iR_\alpha$ where $N_i$ is the density of species $i$ and $R_\alpha$ is the concentration of nutrient $\alpha$. Upon consumption, a fraction of consumed nutrients $1-l$ are assumed to contribute to the growth of microbes and the remaining fraction $l$ is converted to other byproducts. The byproduct conversion flux from nutrient $\alpha$ to nutrient $\beta$ is encoded by $D_{\beta\alpha}$ and thus the matrix $D$ encodes the partitioning of one nutrient to other byproduct nutrients. To conserve fluxes of nutrients, $\sum_\beta D_{\beta\alpha} = 1$ is assumed. Therefore, the total consumption rate of nutrient $\alpha$ by all species is $J^{con}_\alpha = \sum_i J^{con}_{i\alpha} = \sum_i a_{i\alpha}N_iR_\alpha$, while the total production rate of nutrient $\alpha$ is $J^{pro}_\alpha = l\sum_\beta D_{\alpha\beta}J^{con}_\beta = l\sum_\beta\sum_i D_{\alpha\beta}a_{i\beta}N_iR_\beta$. For each species, the growth rate $g_iN_i = \frac{(1-l)\sum_\alpha a_{i\alpha}N_iR_\alpha}{Y}$, which is the sum of consumption rates on all nutrients that are not converted to byproducts divided by the yield $Y$. $g_i$ is also termed as the specific growth rate of species $i$. Overall, the dynamics of the dynamics for concentrations of nutrients $R_\alpha$ and microbial species abundance $N_i$:

$$\frac{dR_\alpha}{dt} = h_\alpha - \delta R_\alpha - J^{con}_\alpha + J^{pro}_\alpha, \tag{S1}$$

$$\frac{dN_i}{dt} = -\delta N_i + g_iN_i. \tag{S2}$$

To obtain the production matrix used in Fig. 5d, we multiply the consumption matrix with the byproduct conversion matrix $D$. More specifically, the production flux of byproduct $\alpha$ by species $i$ is written as $P_{i\alpha} = \sum_\beta D_{\alpha\beta}a_{i\beta}$ which is a sum of all possible byproducts produced by species $i$ when it consumes all available nutrients.

#### Synthetic data from MiCRM

We used the above MiCRM with 10 microbial species and 10 nutrients to generate the synthetic cross-sectional data. All MiCRM parameters such as consumption rates, dilution rates, and byproduct generation rates are assumed to be the same. Different samples are considered as a random sampling of microbial species and nutrients to assemble. Overall, our procedure for generating synthetic data can be divided into three steps: (1) the metapopulation establishment stage: for the metapopulation with all potential species and nutrients, model parameters are drawn from their corresponded probability distributions, (2) the subsampling stage: a fraction of microbial species and nutrients are selected to start the community assembly, and (3) simulation stage: dynamics of sampled species and nutrients are simulated as specified in MiCRM until the synthetic community reaches a steady state. For this community, we collected steady-state relative abundances for all microbial species as the microbial composition, steady-state nutrient concentrations as the corresponding metabolomic profile, and nutrient supply rates as the diet. This 3-stage procedure is repeated many times to create microbial compositions, metabolomic profiles, and diets to form independent samples in the synthetic data.

More specifically, during the first stage, model parameters are determined as follows:

- the chance of one random species consuming one random nutrient is assumed to be 20%. The rate $a_{i\alpha}$ is drawn from the uniform distribution between 0 and 100. After the random drawing, the rate $a_{i\alpha}$ is divided by the number of nutrients the species $i$ can consume to prevent the existence of superbugs.

- the connectance of the byproduct conversion matrix $D$ is assumed to be 50%. In practice, each entry in the matrix $D$ has a probability of 50% to be non-zero. The non-zero entries are drawn from the uniform distribution between 0 and 1. After the drawing, the normalization is imposed to guarantee $\sum_{\beta} D_{\beta\alpha} = 1$.
- the byproduct fraction $l = 0.5$ for all cases.
- the dilution rate $\delta = 0.2$ hour$^{-1}$.
- the yield $Y = 1$.
- the nutrient supply rate $h_\alpha$ is drawn from the uniform distribution between 0 and 1.

During the second stage, for each sample, 50% of species are randomly chosen to be introduced in the initial pool (i.e., $p_{\mathrm{s}} = 0.5$) and nutrients are randomly chosen with the sampling probability $p_{\mathrm{n}}$ to have non-zero supplies (as defined in the nutrient supply rate $h_i$ in the first stage).

#### Modified MiCRM with species-specific byproduct generation and no overlap between consumption and production interactions

The MiCRM above assumes a universal byproduct generation that is encoded by the byproduct conversion matrix $D$ and preserved for all species. This would lead to a case where one metabolite can be consumed and produced by one species at the same time (i.e. overlap between consumption and production interactions for the same metabolite-species pairs). To avoid the overlap, here we considered a more generic case where the byproduct generation is specific to each species. After microbial species $i$ consumes all available nutrients in the community, all consumed nutrients are divided into two parts: (1) a fraction $l$ of total consumed nutrients by species $i$ contributes to the biomass growth, and (2) the other fraction $1 - l$ of total consumed nutrients by species $i$ are converted to byproducts, with their production fluxes proportional to the pre-specified production rate by species $i$ (written as $P_{i\alpha}$). $\sum_{\alpha} P_{i\alpha} = 1$ is imposed to conserve the total flux of nutrients. Other dynamics such as growth dynamics, nutrient consumption, and dilutions are the same as the previous MiCRM. Overall, the dynamics of nutrient concentrations $R_\alpha$ and microbial species abundances $N_i$ are as follows:

$$\frac{dR_\alpha}{dt} = h_\alpha - \delta R_\alpha - \sum_i a_{i\alpha} N_i R_\alpha + \sum_i P_{i\alpha} l \sum_\beta a_{i\beta} N_i R_\beta, \tag{S3}$$

$$\frac{dN_i}{dt} = -\delta N_i + \frac{(1-l)\sum_\alpha a_{i\alpha} N_i R_\alpha}{Y}. \tag{S4}$$

All parameters follow the same definitions specified in the previous MiCRM except for species-specific byproduct production rates $P_{i\alpha}$. The protocol for generating synthetic data by this new version of MiCRM follows the protocol for the previous MiCRM except:

- the chance of one random species producing one random byproduct is assumed to be 50%. $P_{i\alpha}$ is drawn from the uniform distribution between 0 and 1. To avoid the overlap between consumption and production interactions between the same microbe-metabolite pairs, any overlapped entries in the production matrix will be set as zero. After the random drawing and setting values of overlapped entries as zero, the normalization over each species is imposed to guarantee $\sum_\alpha P_{i\alpha} = 1$.

#### Supplementary Figures

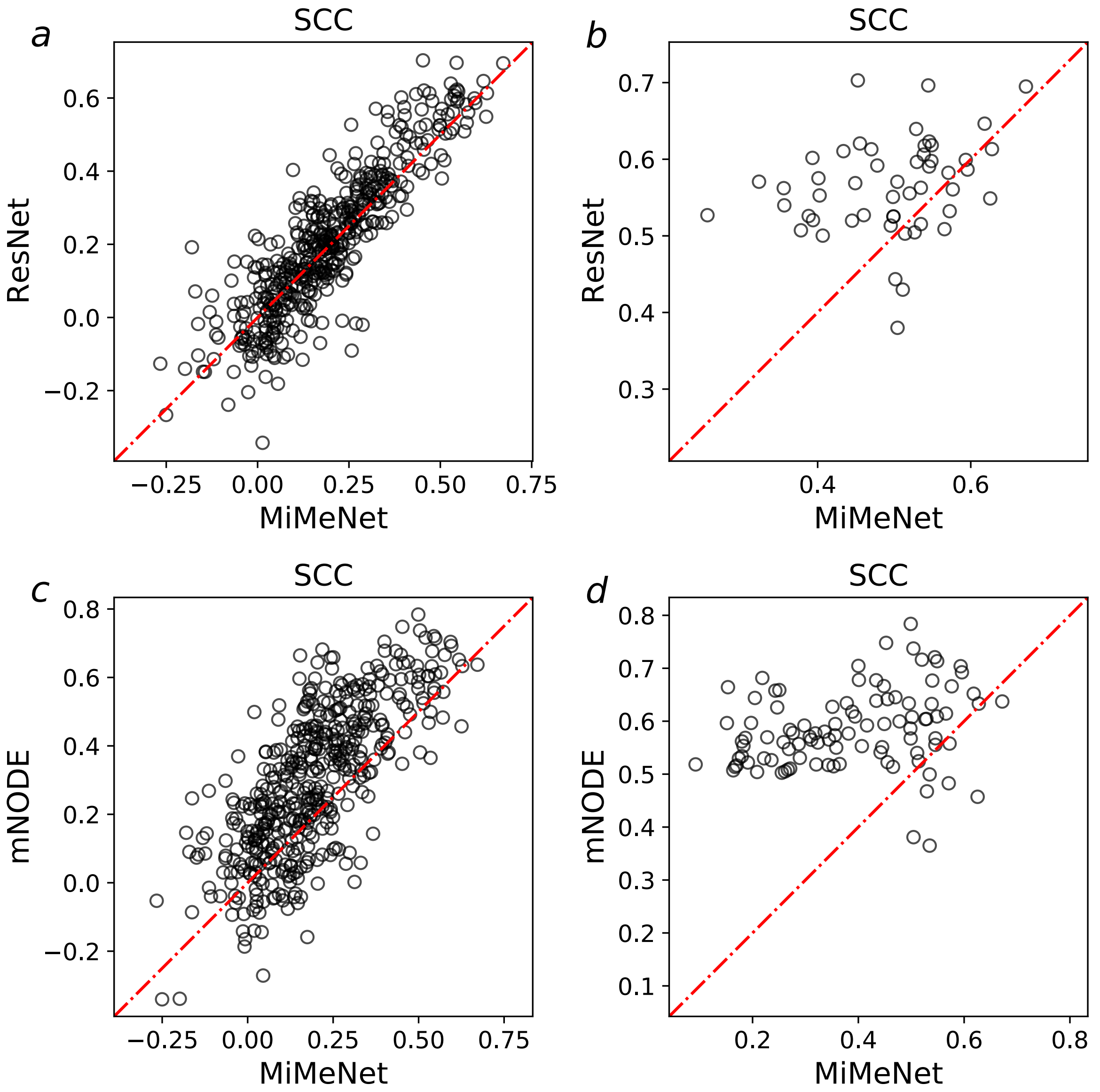


Figure S1: **Comparison of SCCs of annotated metabolites on the test set NLIBD.** For each annotated metabolite, its SCC between its predicted values and true values across samples is computed for all computational methods. **a** Comparison of SCCs of all annotated metabolites between MiMeNet and ResNet. **b** Comparison of SCCs of all well-predicted annotated metabolites between MiMeNet and ResNet. Well-predicted metabolites are metabolites that have SCCs larger than 0.5 according to either MiMeNet or ResNet. **c** Comparison of SCCs of all annotated metabolites between MiMeNet and mNODE. **d** Comparison of SCCs of all well-predicted annotated metabolites between MiMeNet and mNODE. Well-predicted metabolites are metabolites that have SCCs larger than 0.5 according to either MiMeNet or mNODE.

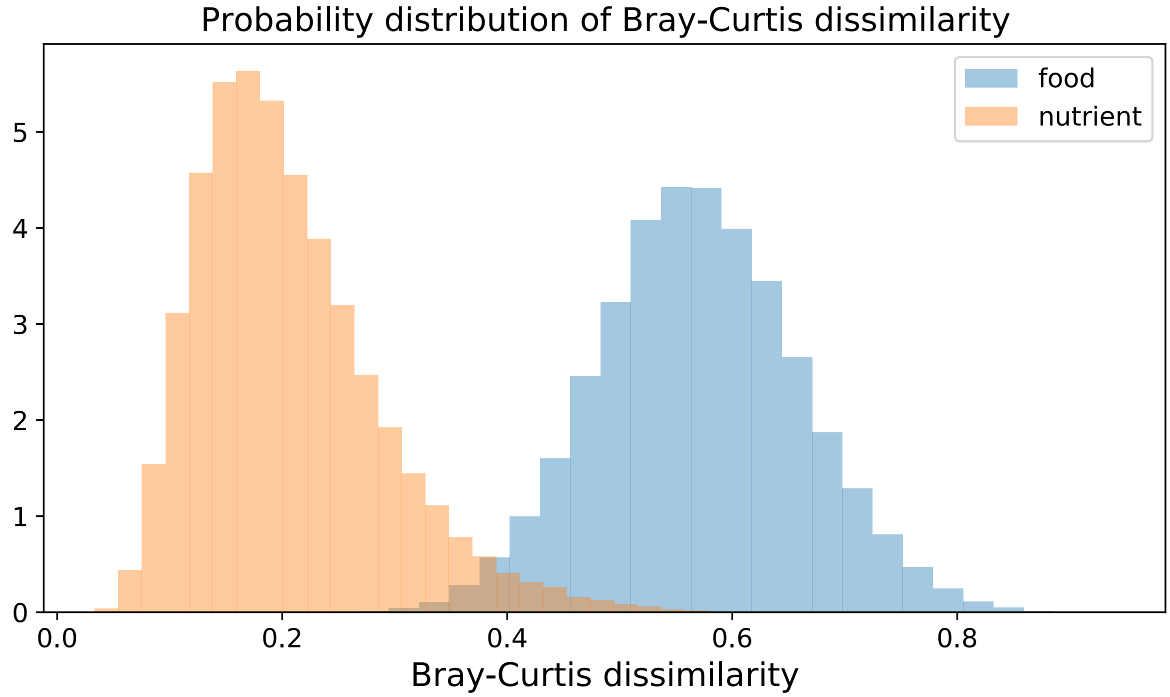


Figure S2: **Bray-Curtis dissimilarity of the food consumption profiles in FFQs and nutritional profiles across samples in VDAART.** The distribution of Bray-Curtis dissimilarity for all paired food consumption profiles in FFQs and nutritional profiles.

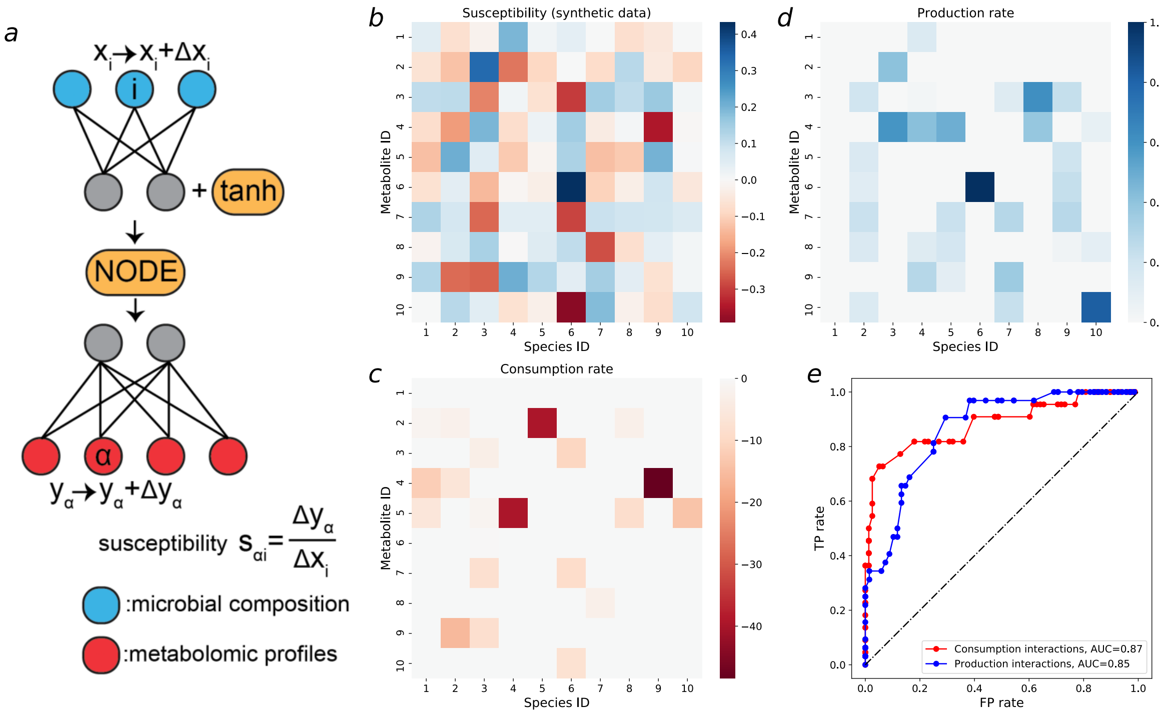


Figure S3: **Using susceptibility of metabolite concentrations to microbial relative abundances of well-trained mNODE to infer microbe-metabolite interactions on new synthetic data.** New synthetic data in this figure are generated by the microbial consumer-resource model with species-specific byproduct generations and without the overlap between consumption and production interactions between microbes and metabolites. See the supplemental information for more details of this model. **a** The susceptibility of the concentration of metabolite $\alpha$ ($y_\alpha$) to the relative abundance of species $i$ ($x_i$), denoted as $s_{\alpha i}$, is defined as the ratio between the deviation in the concentration of metabolite $\alpha$ ($\Delta y_\alpha$) and the perturbation amount in the relative abundance of species $i$ ($\Delta x_i$) **b** Susceptibility values for all microbe-metabolite pairs in the synthetic data. **c** The ground-truth consumption matrix and corresponding rates in synthetic data. All consumption rates are shown as negative values for the convenience of comparison with panel b. **d** The ground-truth production matrix and corresponding rates in synthetic data. **e** The ROC (Receiver Operating Characteristic) curve based on TP rates and FP rates which are obtained by setting different susceptibility thresholds for classifications of interactions.

**Supplementary Data Legends**

**Supplementary Data 1:** Susceptibility values for PRISM + NLIBD.

**Supplementary Data 2:** Susceptibility values for lung samples from patients with cystic fibrosis.

**Supplementary Data 3:** Susceptibility values for soil biocrust samples.

**Supplementary Data 4:** Susceptibility values for fecal samples of children in VDAART.

**Supplementary Data 5:** Susceptibility values for blood plasma samples of children in VDAART.
